# Molecular basis of Cyclin B1-CDK1 recruitment by MAD1

**DOI:** 10.64898/2026.11.01.762963

**Authors:** Mihkel Örd, Natalia de Martin Garrido, Matthew J. Winters, Mythili S. Subbanna, Johanna Kliche, Caroline Benz, Sophie L. Williams, Ylva Ivarsson, Basil J. Greber, Peter M. Pryciak, Norman E. Davey

**Affiliations:** University of Cambridge, CRUK Cambridge Institute, Cambridge, UK; The Institute of Cancer Research, Chester Beatty Laboratories, 237 Fulham Road, London, SW3 6JB, UK; Institute of Bioengineering, University of Tartu, Nooruse 1, 50411 Tartu, Estonia; Department of Biochemistry and Molecular Biotechnology, University of Massachusetts Chan Medical School, Worcester, MA 01605, USA; Department of Chemistry for Life Sciences, Uppsala University, Husargatan 3, 751 23 Uppsala, Sweden; Science for Life Laboratory, Uppsala University, Uppsala, Sweden; Department of Cell and Molecular Biology, St. Jude Children’s Research Hospital, Memphis, TN, USA

**Author notes:** correspondance: Peter M. Pryciak, Norman E. Davey. equal contribution (Natalia de Martin Garrido, Matthew J. Winters).

## Abstract

The interaction between Cyclin B1-CDK1 and MAD1 recruits kinase activity to the nuclear pore complex and kinetochore corona to sustain spindle assembly checkpoint (SAC) signaling. Here, we characterize a short linear motif (SLiM) within MAD1 that mediates its interaction with Cyclin B1. This SLiM peptide is specific for B-type cyclins and lacks the canonical “RxL” cyclin-binding consensus sequence, suggesting a distinct mode of cyclin recognition. Using a combination of deep mutational scanning, structural analysis by cryogenic electron microscopy, and binding affinity measurements, we define the molecular basis of the Cyclin B1-MAD1 interaction. These approaches reveal that a short MAD1 peptide binds as an α-helix to the RxL-binding hydrophobic patch of Cyclin B1. The interaction employs hydrophobic contacts that overlap with those used by canonical RxL motifs, but is distinguished by additional contacts and sequence features that confer preferential recognition by B-type cyclins while excluding other cyclin types. We also show that the MAD1 motif is sufficient to confer B-type cyclin-specific substrate phosphorylation in mitotic cells. Together, these findings reveal a non-canonical, B-type cyclin-specific helical binding mode that ensures selective Cyclin B1-CDK1 recruitment at two critical stages of SAC signaling, and suggest potential strategies for selective therapeutic targeting of this interaction.

## Introduction

Cyclin B1 is a regulatory subunit of Cyclin-Dependent Kinase 1 (CDK1), with which it forms the Cyclin B1-CDK1 complex ^1^. This kinase complex is a central driver of the transition from G2 phase into mitosis, phosphorylating numerous proteins that promote mitotic entry and progression ^2,3^. Cyclin B1 accumulates during S and G2 phases and reaches peak abundance in early mitosis. At the metaphase-to-anaphase transition, Cyclin B1 is targeted for ubiquitin-mediated degradation by the Anaphase-Promoting Complex/Cyclosome (APC/C), thereby promoting mitotic exit ^4,5^. In addition to this temporal regulation, Cyclin B1 is subject to spatial control. During interphase, Cyclin B1 shuttles between the cytoplasm and nucleus but remains predominantly cytoplasmic. Its nuclear accumulation at prophase triggers a switch in localized CDK1 activity, an important regulatory event in mitotic progression ^6,7^.

Cyclin B1 also associates with multiple subcellular structures, including chromatin, the nuclear envelope, and kinetochores ^8^. Association with one or more of these subcellular structures is thought to be important for Cyclin B1-CDK1 to promote signaling of the spindle assembly checkpoint (SAC), which prevents progression to anaphase until all mitotic kinetochores are occupied by spindle microtubules. A central scaffolding protein of the SAC is MAD1, which acts at unattached kinetochores to catalyze the conformational activation of MAD2 and nucleate assembly of the mitotic checkpoint complex (MCC) ^9^. A direct interaction between MAD1 and Cyclin B1 is critical for this process ^10^. During interphase, MAD1 localizes to the nuclear pore complex (NPC) through its interaction with the nucleoporin TPR ^11,12^. Cyclin B1-CDK1 recruited through MAD1 promotes phosphorylation of NPC components and facilitates MAD1 release prior to nuclear envelope breakdown ^13^. Localization of both Cyclin B1 and MAD1 to the kinetochore corona promotes strong SAC signaling, whereas disruption of the MAD1-Cyclin B1 interaction prevents kinetochore localization of Cyclin B1, weakening the SAC, and increasing chromosomal instability ^13,14^.

The interaction between MAD1 and Cyclin B1 was first indicated by their co-immunoprecipitation ^12^, and subsequent work established a direct physical interaction through the N-terminal portion of the MAD1 extended coiled-coil domain ^10^ (Figure 1A). Truncation analyses and the loss of Cyclin B1 binding exhibited by the MAD1β isoform, which naturally lacks exon 4 ^15^, collectively narrowed the binding region to residues 41–62 of MAD1 ^13,14^. Mutagenesis within this region identified three acidic residues, Glu52, Glu53, and Glu56, as critical determinants, with charge-reversal mutations (Glu52Lys, Glu53Lys, and Glu56Lys) abolishing Cyclin B1 binding ^13,14^.

**Figure 1.**
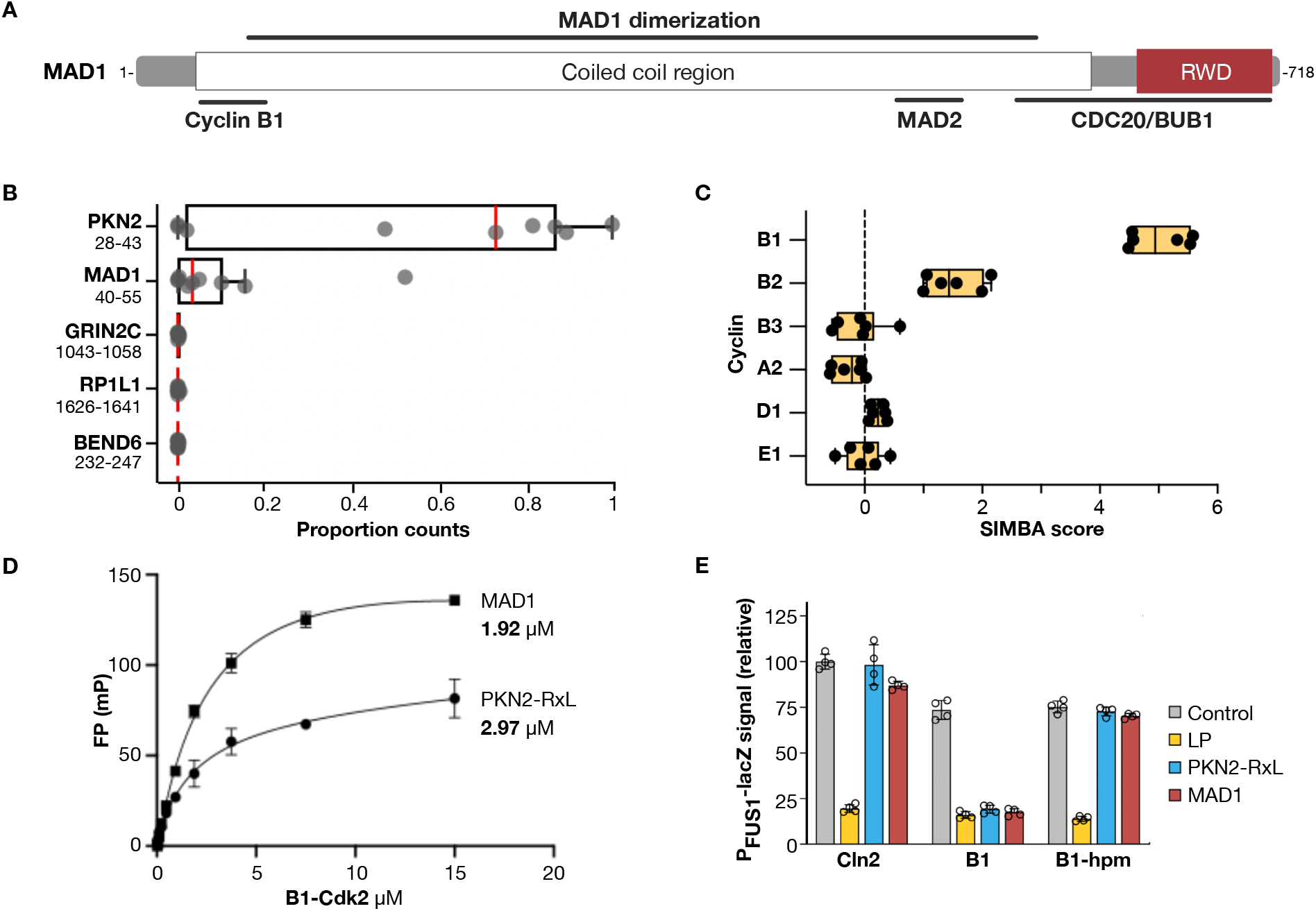
A 16 amino acid peptide from MAD1 is sufficient for specific Cyclin B1 binding. **(A)** Diagram of MAD1 showing domain structure and regions involved in dimerization or binding to Cyclin B1, MAD2, CDC20 and BUB1. RWD: RING finger, WD repeat, DEAD-like helicase domain. **(B)** Proportion of NGS counts of the top five binders for Cyclin B1 in a ProP-PD screen (n=9). **(C)** SIMBA scores for binding of the MAD1 peptide to six Cyclins (n = 6). **(D)** Binding of Cyclin B1 – CDK2 complex to FITC-MAD1 and FITC-PKN2-RxL peptides, measured by fluorescence polarization (mean ± standard deviation, n = 2). **(E)** Suppression of mating signal (*PFUS1-lacZ*) by unfused Cln2, Cln2 fused to wild-type cyclin B1, or Cln2 fused to hydrophobic patch mutant cyclin B1 (B1-hpm). The signaling protein Ste20^Ste5PM^ harbored a control motif, a Cln2-binding motif (LP), an RxL motif (PKN2-RxL), or a MAD1 motif. Mean ± SD (n = 4).

Notably, the Cyclin B1-binding region of MAD1 lacks a canonical RxL cyclin-binding motif ^16,17^, the most prevalent class of cyclin-docking element, raising the possibility that MAD1 engages Cyclin B1 through an alternative binding mechanism. Non-canonical docking mechanisms that engage the canonical cyclin hydrophobic patch through non-RxL sequences have been described in both yeast and human proteins, including motifs specific to the S-phase and mitotic B-type cyclins Clb5 and Clb3 ^18,19^, and a bipartite reverse-orientation engagement of the Cyclin A hydrophobic patch by Skp2 ^20^. Beyond the canonical hydrophobic patch, cyclins also harbor alternative docking surfaces. Cyclin B1 possesses a conserved positively charged phosphate-binding pocket that accommodates a phosphoserine-containing loop of separase ^21^, while D-type cyclins recognize an α-helix in the Rb C-terminus through a surface distinct from the canonical hydrophobic patch ^22^. Whether the MAD1 interaction employs a non-canonical binding mode for the Cyclin B1 hydrophobic patch, engages the phosphate-binding pocket, or exploits an altogether distinct mechanism, has remained unknown.

In this study, we identify a short peptide within MAD1 that is sufficient for binding Cyclin B1 and define the recognition features underlying this interaction. Structural analysis reveals the molecular architecture of the MAD1-Cyclin B1 complex, while saturation mutagenesis identifies residues that determine binding affinity and cyclin selectivity. We further show that this MAD1 motif is sufficient to promote substrate phosphorylation in mitotic cells. Together, these findings define the structural and mechanistic basis by which a non-canonical, B-type cyclin-specific helical motif in MAD1 ensures the selective recruitment of Cyclin B1-CDK1 at critical stages of SAC signaling.

## Results

### Defining a MAD1 region sufficient for binding Cyclin B1

To identify peptide sequences capable of binding Cyclin B1, we used Proteomic Peptide Phage Display (ProP-PD) to screen Cyclin B1 against a library of approximately one million peptides derived from the predicted disordered regions of the human proteome, tiled as 16-mer peptides overlapping by 12 amino acids ^23^. Two predominant Cyclin B1-binding peptides emerged from this screen: one derived from serine/threonine-protein kinase N2 (PKN2), and the other from MAD1 (Figure 1B). We previously showed that the PKN2 peptide contains an atypical RxL-like cyclin-binding motif that is bound by several human cyclins ^17^. The MAD1 peptide found in the ProP-PD selection spans residues 40 to 55 (SLQMQYQQSMQLEERA), which overlaps the region previously implicated in Cyclin B1 binding ^13,14^. This region includes conserved acidic residues previously shown to contribute to the MAD1-Cyclin B1 interaction, including Glu52 and Glu53, and lies immediately adjacent to Glu56 ^13^. Thus, the peptide identified by ProP-PD defines a short region of MAD1 that is sufficient for interaction with Cyclin B1.

### MAD1 is a Cyclin B-specific hydrophobic patch binder

To determine the cyclin-binding specificity of the MAD1 peptide, we used Systematic Intracellular Motif Binding Analysis (SIMBA) ^17,24^, a quantitative yeast-based peptide binding assay, to test multiple human cyclins. The assay revealed a striking preference of the MAD1 peptide for B-type cyclins, with negligible binding to other cyclin classes (Figure 1C). Furthermore, the MAD1 peptide bound both Cyclin B1 and Cyclin B2 but not Cyclin B3. Binding to Cyclin B2 was weaker than binding to Cyclin B1, suggesting that this interaction may functionally distinguish between these closely related B-type cyclins. We quantified the strength of this interaction using fluorescence polarization (FP) (Figure 1D). The MAD1 peptide bound Cyclin B1 with a Kd of 1.9 µM, an affinity similar to that of the strong-binding RxL peptide derived from PKN2 (3.0 µM).

We next sought to define the Cyclin B1 surface recognized by MAD1. Because canonical RxL motifs interact with the cyclin hydrophobic patch ^25^, we initially tested MAD1 binding to Cyclin B1 proteins containing mutations in this surface. These mutations markedly reduced MAD1 binding in SIMBA assays (Figure 1E), indicating that the hydrophobic patch forms the principal docking surface for MAD1. Together, these experiments establish a 16-residue MAD1 peptide as a *bona fide* docking element specific for the hydrophobic patch of B-type cyclins.

### MAD1 binds the Cyclin B1 hydrophobic patch as an α-helix

To investigate how MAD1 engages the Cyclin B1 hydrophobic patch despite lacking a canonical RxL motif, we determined the structure of the Cyclin B1-CDK1 complex bound to a 16-residue peptide derived from MAD1 by cryogenic electron microscopy (cryo-EM) (Figure 2A). Data collection and processing yielded a map at 3.3 Å resolution (Supplementary Figure S1, Supplementary Fig. S2A-D, Supplementary Table S1). The Cyclin B1-CDK1 complex adopted an overall fold consistent with previously determined structures of Cyclin-CDK holoenzymes ^26,27^. In addition, well-defined density corresponding to a 14 amino acid α-helix was observed for MAD1 residues 40-53 (Figure 2B; Supplementary Figure S2E). The MAD1 peptide buried approximately 600 Å² of surface area upon binding Cyclin B1, with Leu41, Tyr45, and Met49 making particularly prominent contributions to the interface (Figure 2C). The peptide occupied the canonical RxL-binding hydrophobic patch, which is formed by residues contributed by helices α1, α3, and α4, including the MRAIL-containing α1 helix. The MAD1 helix was oriented in the same N-to-C direction as typical RxL-containing peptides. Strikingly, MAD1 contributes three hydrophobic cyclin-contacting residues that occupy positions corresponding to the previously described Φ_N_, Φ_0_, and Φ_C_ positions of RxL motifs ^17^ (Figure 2D), despite using noncanonical residues at each position. In RxL motifs, the Φ_N_, Φ_0_, and Φ_C_ positions are typically Ala/Val/Cys, Leu, and Leu/Phe, respectively ^17^. In the MAD1 helix, Leu41 occupies the Φ_N_ position, Tyr45 the Φ_0_ position, and Met49 the Φ_C_ position. In contrast to canonical RxL motifs, MAD1 lacks the P_-2_ basic residue corresponding to the “R” of the RxL motif. This is consistent with prior findings that B-type cyclins show a reduced dependence on a basic P_-2_ residue ^17^ due to their lack of a complementary acidic surface found in other cyclin classes ^16^.

**Figure 2.**
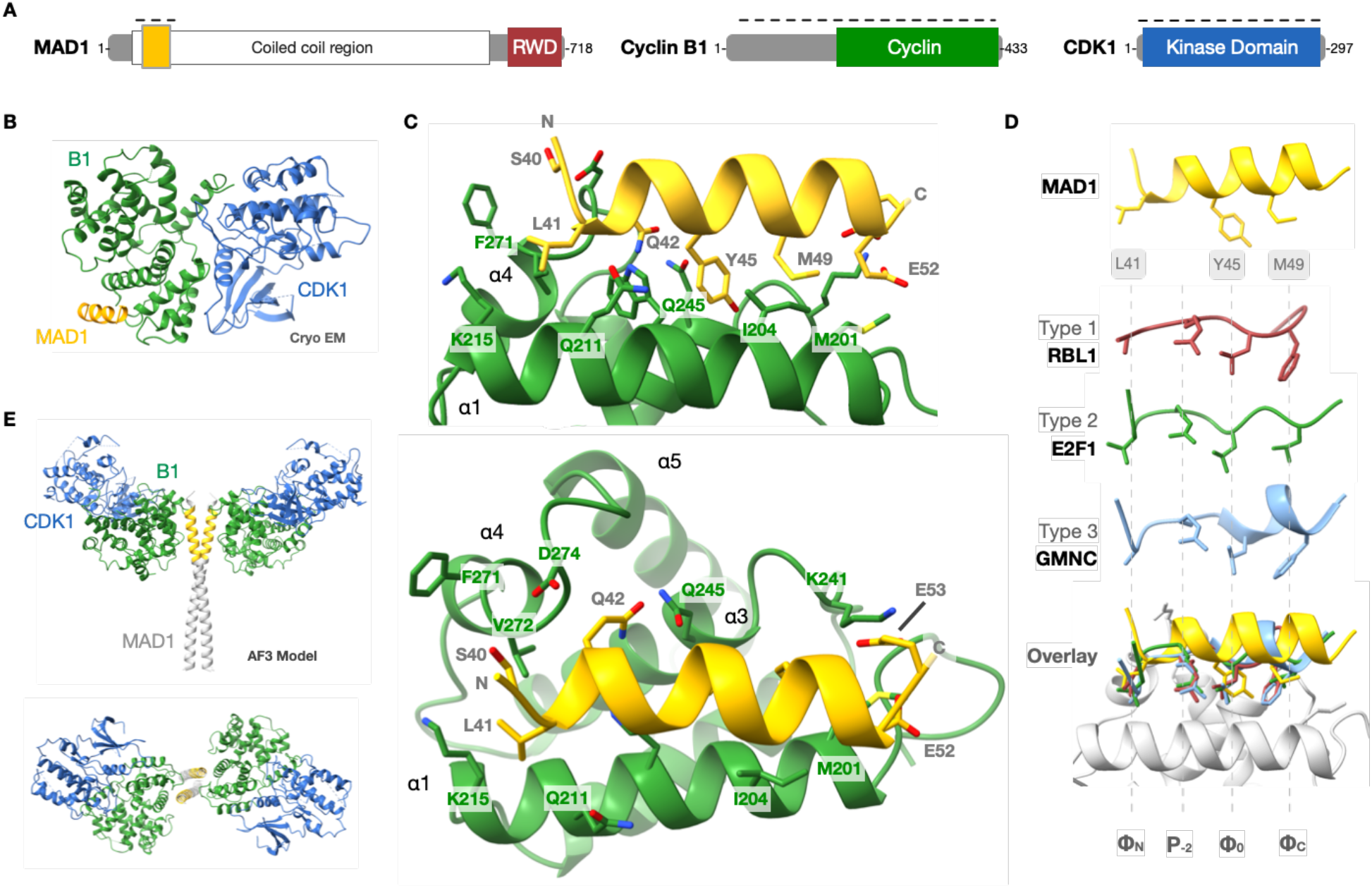
The MAD1 helix binds the hydrophobic patch of Cyclin B1. **(A)** Domain architecture of MAD1, Cyclin B1 and CDK1. **(B)** Cryo-EM structure of the Cyclin B1 - CDK1 - MAD1 complex. **(C)** Detailed view of the bound helix and binding surface. **(D)** Comparison of the MAD1 helix to type 1, 2, and 3 RxL peptides. **(E)** AlphaFold3 (AF3) model of the N-terminal region of the MAD1 coiled-coil with the cryo-EM structure of the Cyclin B1 - CDK1 - MAD1 complex superimposed.

The MAD1 helix also forms contacts that are not generally observed in RxL-cyclin interactions. MAD1 Gln42 buries approximately 111 Å² of surface area and contacts Cyclin B1 residues Thr276 and Gln245, while toward the C terminus of the helix, MAD1 Glu53 interacts with Cyclin B1 Lys241 (Figure 2C). In contrast, MAD1 residues on the distal face of the helix (Met43, Gln44, Gln46, Gln47, and Gln50) make little or no direct contact with the Cyclin B1 surface. Additionally, comparison of peptide-bound and apo Cyclin B1 structures ^26,27^ revealed a small conformational shift of residues 271-273 at the C-terminus of the α4 helix to accommodate MAD1 binding (Supplementary Figure S2F). In cells, MAD1 exists as a constitutive dimer ^28,29^, and AlphaFold3 modeling ^30^ suggests that its coiled-coil could simultaneously accommodate two Cyclin B1 molecules in a back-to-back arrangement (Figure 2E). Together, the structural data reveal that MAD1 uses an α-helical binding mode to occupy the canonical cyclin hydrophobic patch while making a distinctive set of molecular contacts.

### The MAD1 helix has strict binding determinants

To systematically define the sequence determinants of MAD1 recognition by Cyclin B1, we performed deep mutational scanning (DMS) of the MAD1 peptide (residues 40-55) using SIMBA ^24^. The results were broadly consistent with the cryo-EM structure and provided a comprehensive view of the amino acids tolerated at each position of the motif (Figure 3A-C). Position 40 strongly favored Thr or Ser, whereas position 41 showed a stringent preference for Leu, with Met also tolerated. Position 42 required His, Asn, or Gln, while position 45 strongly favored an aromatic residue, with Phe, Trp, and Tyr all supporting binding. At position 49, Met was strongly preferred, although other aliphatic residues were tolerated, indicating some flexibility within this portion of the hydrophobic-binding pocket. Positions 51-53 favored acidic residues, whereas basic residues were disfavored, suggesting an important contribution from charge complementarity. Interestingly, however, preferences at the two native acidic positions differed: while Glu and Asp were equally preferred at residue 53, aliphatic substitutions were preferred over Asp at residue 52, consistent with the structural observation that Glu52 abuts a nonpolar surface of cyclin B1 whereas Glu53 directly faces an electropositive surface. A further constraint evident throughout the MAD1 motif was that Pro and Gly substitutions were generally poorly tolerated. This pattern is consistent with a requirement for MAD1 to adopt the observed α-helical conformation when bound to Cyclin B1. By contrast, positions corresponding to Met43, Gln44, Gln46, Gln47, and Gln50, which make little or no contact with the Cyclin B1 surface in the cryo-EM structure, tolerated a broad range of substitutions other than helix disruptors (Figure 3B). Thus, the mutational landscape closely reflects the binding mode observed structurally, and it defines both side-chain-specific interactions and a broader requirement for α-helical propensity.

**Figure 3.**
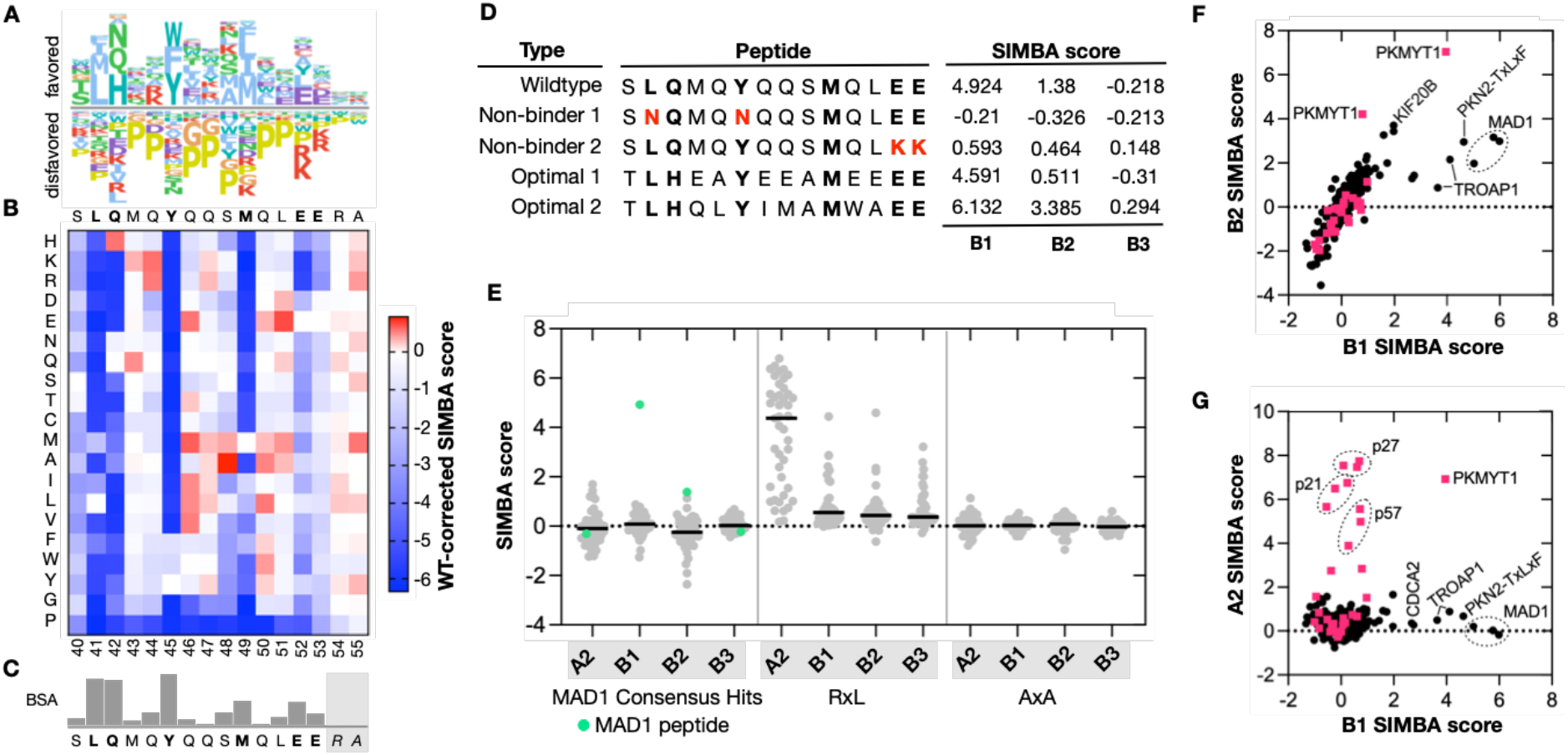
Deep investigation of the Cyclin B1 - MAD1 interaction affinity and specificity determinants. **(A)** Sequence logo summarizing SIMBA deep mutational scanning (DMS) of the MAD1 peptide against Cyclin B1. **(B)** Per-substitution heatmap of the same DMS data. **(C)** Buried surface area (BSA) per peptide residue of the MAD1 peptide in the Cyclin B1–MAD1 complex. Residues 54-55 lack BSA values because they were not resolved in the structure. **(D)** Peptide variants designed from the DMS data to increase (Optimal) or abolish (Non-binder) Cyclin B1 binding. **(E)** Cyclin binding measurements for 43 human peptides that contain the MAD1 cyclin B binding motif consensus sequence, together with RxL peptides and their corresponding AxA negative control peptides from a high-throughput SIMBA experiment. **(F-G)** Comparison of Cyclin B1/B2 (panel F) and Cyclin B1/A2 (panel G) binding to 16-mer predicted hydrophobic patch interacting peptides extracted from Cyclin B1 and B2 Predictome complexes. Pink squares, RxL motifs; black circles, non-RxL motifs. Data is from a high-throughput SIMBA experiment.

### Optimizing Cyclin B1 binding

The sequence preferences revealed by DMS provided a framework for rationally manipulating the affinity of the MAD1 motif. Because the DMS screen quantified the relative preference for every possible residue at each position, we could select preferred amino acids at each site to maximize binding affinity. Notably, the wild-type residue was already optimal at each contact position except Gln42, where substitution with His was favored (Figure 3A-B). We therefore combined Gln42His with additional favorable substitutions at non-contact positions to design optimized binder peptides (Figure 3D). One of these two optimized sequences, TLHQLYIMAMWAEE, bound more strongly than the wild-type MAD1 peptide to both Cyclins B1 and B2. By contrast, two nonbinding control peptides, which incorporated substitutions at key contact positions identified by DMS, showed no detectable binding to Cyclin B1, confirming that these positions are indeed required for the interaction. These results demonstrate that the DMS-derived sequence landscape can be used to engineer the affinity and cyclin specificity of the MAD1 helical motif.

### The MAD1 helix represents an uncommon Cyclin B1-binding mode

Given the widespread occurrence of RxL cyclin-binding motifs ^17^, we asked whether MAD1-like helical motifs might also be present in other human proteins. Based on the specificity determinants obtained by DMS, we defined a loose MAD1-like consensus sequence, [^KRL]L[HNQ][^P][^P][FWY][^PG][^PG][HKRDENQSCMAL][MIFL][^P][^P][^RKWGP][^RK]. Using SLiMSearch ^31^, we identified 43 peptides matching this consensus within predicted disordered regions of the human proteome. However, none of these peptides displayed detectable binding to Cyclin B1 when tested by SIMBA (Figure 3E, Supplementary Table S2), suggesting that the sequence requirements for MAD1-like recognition cannot be fully captured by a simple linear consensus.

As an independent strategy, we searched the Predictomes database, which contains AlphaFold-predicted structure models of protein-protein interactions ^32^, for complexes in which a linear peptide was predicted to engage the hydrophobic patch of Cyclin B1 or Cyclin B2. Of 327 16-mer peptides subsequently tested by SIMBA, the great majority showed no detectable binding (Figure 3F, Supplementary Table S3). Cyclin B1 binding was detected for three additional forms of the MAD1 peptide that overlap the original 16-mer (GSLQMQYQQSMQLEER, PGSLQMQYQQSMQLEE, and APGSLQMQYQQSMQLE), as well as two previously characterized RxL-type peptides derived from PKMYT1 (FEPRNLLSLFEDTLDP) and PKN2 (SDSTVTKLDFDLEPEP), while cyclin A2 recognized exclusively canonical RxL peptides (Figure 3G). Five additional noncanonical peptides displaying weak SIMBA interactions were tested for their ability to capture Cyclin B1 or Cyclin B2 from cell lysates. Of these, only a peptide derived from KIF20B (INFDGIKLDLSHEFSL) showed detectable binding in the pulldown assay (Supplementary Figure S3A). Interestingly, this peptide preferentially bound Cyclin B2 in both SIMBA and pulldown experiments.

As a final approach to find additional similar motifs, we examined MAD1 peptides from eight additional vertebrate species, each differing from the human sequence at 4 to 10 positions. The mouse MAD1 ortholog, which differs only at four non-core positions within the 16-mer, showed strong binding to both Cyclin B1 and Cyclin B2 (Supplementary Figure S3B, Supplementary Table S3). Peptides from more distantly related vertebrates contain one or more core positions substituted with largely physicochemically similar residues. Nevertheless, most bound Cyclin B1 weakly, except for those from the frog *Xenopus tropicalis* and the seahorse *Hippocampus comes* (Supplementary Figure S3B, Supplementary Table S3), of which the latter has only one of the core positions changed to its second-favored residue (Glu52Leu). The findings above collectively suggest that the MAD1 helix represents a mode of cyclin binding that is highly uncommon, and possibly unique within the human proteome, which likely reflects stringent requirements at multiple contact positions across the peptide sequence.

### Cyclin pocket differences dictate MAD1 selectivity

The specificity of the MAD1 peptide for B-type cyclins is remarkable, given that it binds to the same hydrophobic patch used by multiple types of cyclins to bind RxL motifs. In particular, Cyclin A2 binds to the broadest array of RxL motifs ^17^, yet it does not bind MAD1. Notably, this specificity is not captured by structure prediction, as AlphaFold3 models the MAD1 peptide bound to the hydrophobic patch of both cyclins with comparable confidence (Supplementary Figure S4A). Therefore, we investigated which features of the cyclin B1 hydrophobic patch confer its preferential recognition by the MAD1 peptide.

Of the 16 Cyclin B1 residues that contact the MAD1 helix (defined as > 5 Å² buried surface area), 10 are identical in Cyclin A2. These include the Cyclin A2 hydrophobic core residues Ile213 and Trp217, as well as polar residues such as Gln254 and Asp283 (Figure 4A, Supplementary Figure S4B). Three of the remaining residues differ between the two cyclins and cluster within or adjacent to the MRAIL-containing α1 helix. However, replacing these Cyclin A2 residues with their Cyclin B1 counterparts (Ser209Asn, Glu220Gln, and Glu224Lys) was insufficient to confer binding to the MAD1 peptide (Figure 4B-D). We therefore tested substitutions throughout a larger interaction surface containing 10 residues that differ between the two cyclins. While small subsets of substitutions did not suffice, MAD1 binding became increasingly detectable as more substitutions were introduced into Cyclin A2, and was greatest when all ten positions were replaced (cycA2*-hp10*) (Figure 4B-D). Notably, this gain of MAD1 binding was accompanied by reduced binding to canonical RxL motif peptides from p27 and E2F1, consistent with the loss of electronegative contacts for basic RxL side chains (Figure 4B, Supplementary Figure S4B). A reciprocal experiment produced the reverse effect: replacing five positions in the Cyclin B1 pocket with the corresponding Cyclin A2 residues (cycB1*-hp5*) reduced MAD1 binding while enhancing interactions with RxL peptides (Figure 4B-D).

**Figure 4.**
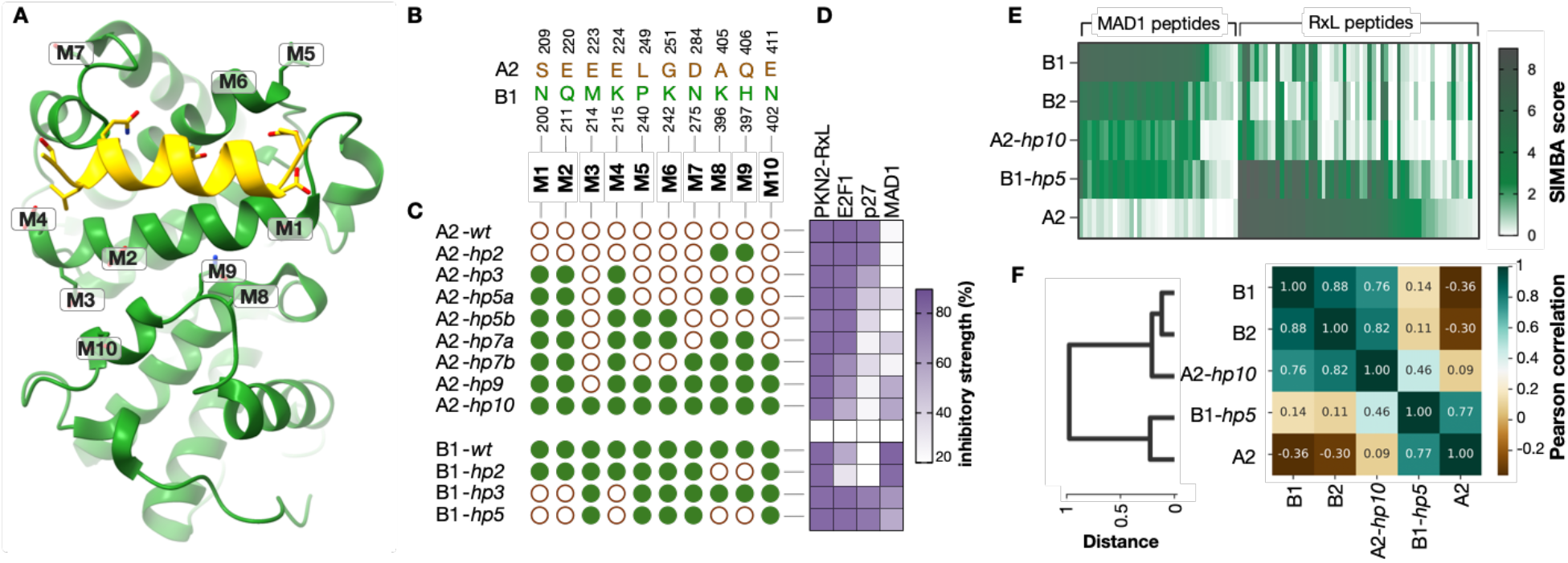
Analysis of cyclin pocket residues controlling MAD1 and RxL motif binding. **(A)** Structure of the Cyclin B1 (green) - MAD1 (yellow) complex overlaid with positions (M1–M10) mutated in the domain pocket swap analysis. **(B)** Amino acids and amino acid positions in Cyclin A2 and B1 mutated during the pocket swap analysis. Labelled M1–M10 as in panel A. **(C)** Mutation groupings tested in the pocket swap analysis. For A2, filled green circles signify that the position was switched to the B1 residue. For B1, empty brown circles signify that the position was switched to the A2 residue. **(D)** Low-throughput SIMBA measurements of peptide binding strength for the pocket swap derivatives in panel C. Inhibition of mating signaling (*PFUS1-lacZ*) for each peptide was calculated as the percent reduction relative to a control peptide (mean; n = 2-3). **(E)** High-throughput SIMBA scores show how cyclin pocket swaps alter binding to MAD1 and RxL peptides. RxL peptides include 51 wild-type and 7 mutated sequences. MAD1 peptides include human MAD1, 8 non-human orthologs, 4 non-binder and optimal variants from Fig 3D, and 25 high-scoring DMS variants. **(F)** Pairwise correlations of cyclin derivatives, using the data in panel E.

Analysis across a larger panel of 58 RxL peptides and 38 MAD1 variants revealed a broad exchange of binding preferences (Figure 4E-F, Supplementary Table S3). Wild-type Cyclin A2 bound a wide range of RxL peptides, of which only a minority bind to Cyclins B1 and B2. The cycA2-*hp10* mutant adopted a peptide-binding profile resembling that of Cyclins B1 and B2, in that it gained binding to MAD1 peptides and lost binding to many RxL peptides, while retaining binding to most of the B1/B2-recognized subset. Conversely, the cycB1-*hp5* mutant shifted toward the binding profile of Cyclin A2, in that it showed reduced binding to MAD1 peptides but increased binding to many RxL peptides. On the peptide side, DMS results showed that no single substitution in the MAD1 peptide was sufficient to confer strong Cyclin A2 binding (Supplementary Figure S4C-D). Furthermore, the MAD1 DMS profiles for all four cyclin variants with detectable MAD1 binding (B1, B2, A2*-hp10*, and B1*-hp5*) showed identical preferences at the core hydrophobic positions (Φ_N_, Φ_0_, and Φ_C_), but differed at positions 44, 48, and 52 (Supplementary Figure S4C, S4E), suggesting that pocket composition subtly shapes the recognition optimum. Together, these results indicate that the distinct peptide preferences of Cyclins A2 and B1 arise from multiple, broadly distributed sequence differences across the cyclin binding pocket.

### The MAD1 helix can encode B-type cyclin-specific mitotic phosphorylation

Finally, we asked if the MAD1 helix could serve as a docking element to confer Cyclin B-dependent substrate phosphorylation in cells. For this, we incorporated the MAD1 helix into a PKMYT1-based phosphorylation reporter (Figure 5A) ^17^. As a first step, we conducted pulldown experiments to confirm that the MAD1 peptide conferred selective binding of the reporter to B-type cyclins in human cell lysates, in contrast to an RxL peptide that binds both A- and B-type cyclins (Figure 5B). Next, we examined reporter phosphorylation during the cell cycle. In cells arrested at prometaphase, when Cyclin B1-CDK1 activity is high, reporters containing the MAD1 docking element showed pronounced phosphorylation-dependent mobility shifts when resolved by Phos-tag gel electrophoresis (Figure 5C). We next monitored phosphorylation after releasing cells from palbociclib-induced G1 arrest (Figure 5D). Reporter phosphorylation closely correlated with the accumulation of Cyclin B1 and was detectably elevated only at the time point corresponding to mitosis (Figure 5E). Thus, the MAD1 helix is not only sufficient for selective Cyclin B binding but can also confer cell-cycle-specific phosphorylation during the period of maximal Cyclin B1-CDK1 activity.

**Figure 5.**
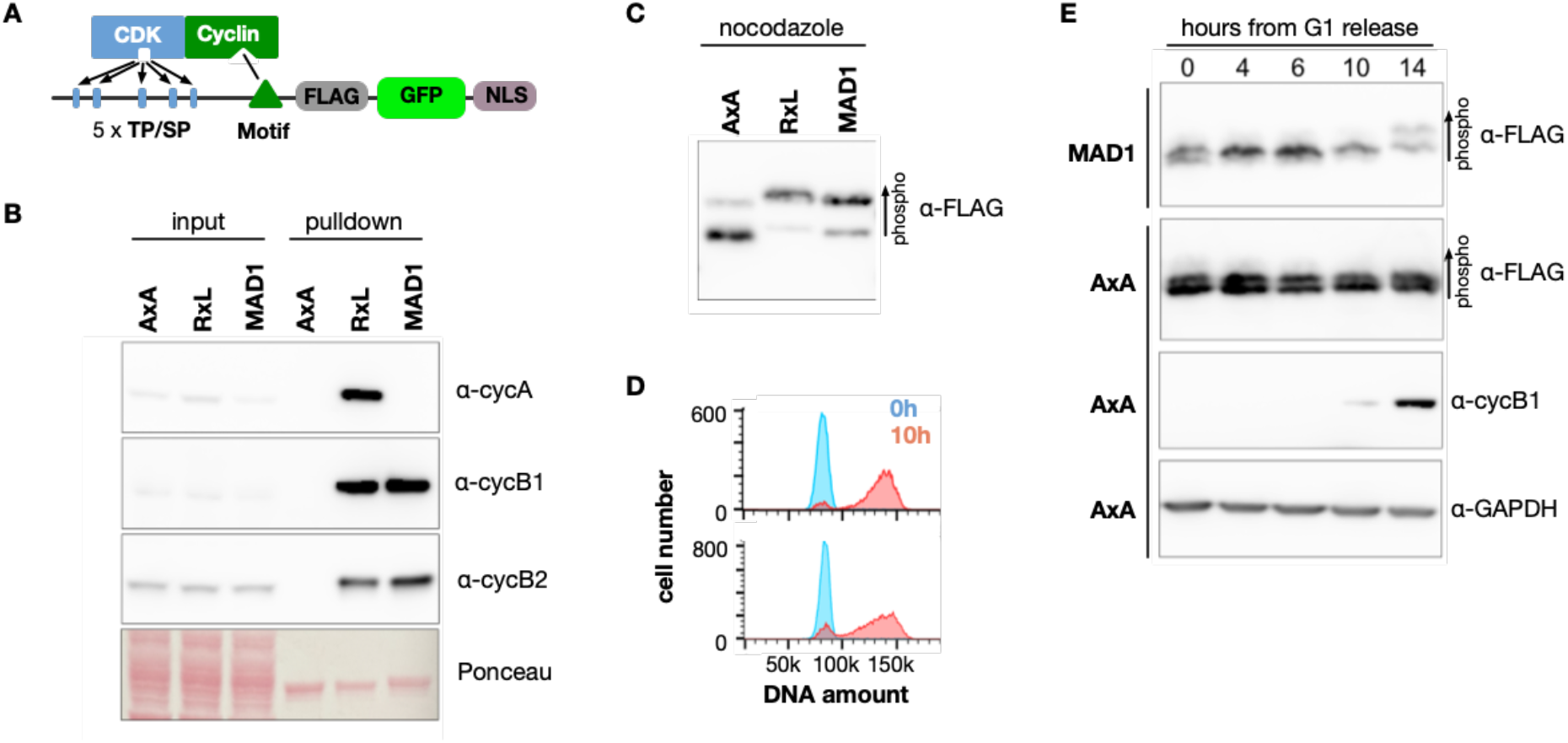
**(A)** PKMYT1-based phosphorylation reporter to test whether inserted motifs recruit cyclin-CDKs. **(B)** Pulldown assays testing reporters with different docking motifs (AxA, RxL, MAD1) for binding to cyclins in human cell lysates. **(C)** Phosphorylation of PKMYT1-based reporters in nocodazole arrested cells. **(D)** DNA content histograms showing cell cycle progression from a synchronized G1 population (0h, blue) to 10h post-release (10h, red). **(E)** Phosphorylation of PKMYT1-based reporters in palbociclib-synchronized cells monitored by Phos-tag SDS-PAGE.

## Discussion

Accurate chromosome segregation requires precise spatial and temporal coordination of CDK1 activity during mitosis ^2^. The spindle assembly checkpoint provides a surveillance mechanism that delays mitotic progression until all chromosomes have established appropriate attachments to the mitotic spindle ^33,34^. An important component of this regulation is the interaction between MAD1 and Cyclin B1-CDK1, which promotes the recruitment of CDK1 activity to nuclear pore complexes and kinetochores ^10,13,14^, and depends on the N-terminal region of MAD1 ^10,13,14^. Here, we defined the molecular interface responsible for this interaction and revealed an unconventional cyclin-binding mechanism that enables selective recruitment of Cyclin B1-CDK1.

Canonical cyclin-binding motifs, most notably the RxL motif, are well-established mediators of cyclin-CDK recognition ^25,35^. We identified a distinct SLiM peptide within the MAD1 N terminus that engages Cyclin B1 despite lacking the defining sequence features of an RxL motif. By combining DMS, cryo-EM structural analysis, and biochemical binding measurements, we define both the structure of this interaction and the sequence determinants underlying its specificity. While our experiments validated the previously identified involvement of MAD1 Glu52 and Glu53 in cyclin B1 binding ^13,14^, they revealed a more extensive interface dominated by nonpolar contacts. The MAD1 peptide bound Cyclin B1 as an amphipathic α-helix in which three hydrophobic side chains contact the canonical cyclin hydrophobic patch. These side chains occupy positions analogous to the hydrophobic Φ_N_, Φ_0_, and Φ_C_ contacts of RxL motifs, but the motif uses noncanonical side chains and lacks the archetypal basic residue that defines RxL peptides. The interaction therefore illustrates how a single cyclin surface can recognize structurally and compositionally distinct docking motifs.

The MAD1 motif extends the range of peptide architectures known to engage the hydrophobic patches of human and yeast cyclins and reinforces the emerging view that the recognition capabilities of this pocket are considerably more permissive than originally appreciated ^17–20,36,37^. MAD1 could in principle have evolved a conventional B1-specific RxL motif to perform the same function, yet the helical motif has notably distinct biochemical properties. In particular, it binds Cyclin B1 more strongly than any of the 50 wild-type RxL peptides we examined while displaying negligible binding to non-B type cyclins. Thus, the helical binding mode combines high-affinity recognition of Cyclin B1 with unusually stringent cyclin selectivity.

Our mutational analysis further demonstrates that this selectivity arises from distributed features of the cyclin hydrophobic patch rather than from a single discriminatory contact. Despite the overall similarity of the Cyclin A2 and Cyclin B1 docking surfaces, replacing a small number of candidate specificity residues was insufficient to confer MAD1 recognition on Cyclin A2. Instead, multiple substitutions across an extended region of the hydrophobic patch were required to switch binding behavior. Importantly, these substitutions not only altered MAD1 recognition but also reciprocally affected binding to canonical RxL peptides. The hydrophobic patches of different cyclins therefore encode distinct recognition landscapes through the collective effects of multiple pocket residues. This provides a molecular explanation for how closely related cyclins can use homologous docking surfaces yet recognize different subsets of peptide motifs. Interestingly, current structure prediction methods do not capture these differences. AlphaFold3 docks the MAD1 peptide onto both Cyclin A2 and B1 with similar configuration and confidence, yet we measure no binding to Cyclin A2. Therefore, while these algorithms can propose plausible docking modes, they do not yet resolve the fine-grained determinants of affinity and specificity for pocket-peptide interactions.

The apparent rarity of MAD1-like sequences further emphasizes the unusual nature of this interaction. Neither sequence-based nor structure-guided searches identified other comparable Cyclin B1-binding peptides. These observations suggest that the helical MAD1 motif is a specialized solution to meet the regulatory requirements of this particular cyclin interaction. The same DMS landscape that explains why such sequences are uncommon also provides a blueprint to engineer MAD1-derived peptides with increased affinity. Interestingly, optimizing the sequence for Cyclin B1 binding simultaneously increased recognition by Cyclin B2, illustrating a potential trade-off between binding strength and specificity within the B-type cyclin family.

Cyclins have attracted increasing interest as drug targets ^38–40^. The recent demonstration that macrocyclic RxL peptide inhibitors can block the hydrophobic patch of both Cyclin A and Cyclin B, selectively killing E2F-high cancer cells and showing anti-tumor activity in patient-derived xenograft models, establishes that this surface is pharmacologically tractable ^38^. Understanding the molecular specificity of the MAD1-Cyclin B1 interaction may provide a route toward selective therapeutic manipulation of Cyclin B1-CDK1 function. Molecules or peptides that disrupt this SLiM-mediated interaction could, in principle, enable selective inhibition of Cyclin B1-CDK1 recruitment to the kinetochore or NPC, without affecting the broader, essential functions of the Cyclin B1-CDK1 complex or RxL-mediated recruitment by the extended Cyclin family. This could enhance the efficacy of anti-mitotic therapies, which are often limited by toxicity and resistance, by specifically destabilizing the spindle assembly checkpoint in cancer cells that are hyper-dependent on these pathways for chromosomal stability. Finally, the molecular definition of this interface provides tools for addressing the biological functions of the MAD1-Cyclin B1 interaction, which remain incompletely resolved. Mutations that selectively disrupt MAD1 binding while preserving interactions with RxL-dependent Cyclin B1 partners should enable direct analysis of the functional contribution of the MAD1-Cyclin B1 interaction to cell division and to the recognition of CDK substrates at kinetochores and NPCs. Conversely, cyclin pocket mutations that shift preferences between RxL and MAD1 binding provide an opportunity to determine whether restricting MAD1 recognition to B-type rather than A-type cyclins has a critical regulatory benefit, such as controlling the timing of specific mitotic events. More broadly, these findings illustrate how diversification of cyclin-peptide recognition can generate functional distinctions among closely related cyclin-CDK complexes and their numerous interaction partners.

## Acknowledgements

We thank Danny D’Amore for critical feedback on the manuscript, Chris Richardson for support with high-performance computing, and Cancer Research UK Cambridge Institute Research Instrumentation and Cell Services for cell line testing. We acknowledge the London Cryo-EM Consortium (LonCEM), which is supported by Wellcome grant 206175/Z/17/Z, for Titan Krios access. We thank Nora Cronin for support of data collection. N.E.D. and M.O. were funded by a Cancer Research UK Senior Cancer Research Fellowship (C68484/A28159). N.E.D. was supported by the American Lebanese Syrian Associated Charities (ALSAC). M.O. was supported by Cancer Research UK SEBINT-2024/100003 and Estonian Research Council research grant PRG2149. Work by M.J.W, M.S.S, and P.M.P. was funded by grants from the NIH to P.M.P. (R01GM145795 and R35GM163698). B.J.G. was supported by a Career Development Award from the Medical Research Council of the UK, grant MR/V009354/1. Work by J.K. and C.B. was funded by grants for the Swedish Research Council to Y.I. (2020-03380).

## Author contributions

**M.J.W., M.S.S., N.M.G., V.I.C., J.K., C.B:** Investigation, Data analysis. **S.L.W.:** Resources. **Y.I.:** Supervision, Funding acquisition, Writing - Review & Editing. **B.J.G.:** Supervision, Visualization, Data analysis, Funding acquisition, Writing - Review & Editing. **M.O.:** Conceptualization, Investigation, Visualization, Data analysis, Writing – original draft. **P.M.P.:** Conceptualization, Investigation, Project administration, Supervision, Visualization, Data analysis, Funding acquisition, Writing – original draft. **N.E.D.:** Conceptualization, Project administration, Supervision, Visualization, Data analysis, Funding acquisition, Writing – original draft.

## Competing interests

N. E. D. is a scientific co-founder of RIME therapeutics. The rest of the authors declare that they have no competing interests.

## Data availability

The raw data generated in this study is available in the supplementary material. The atomic models and cryo-EM maps of the CDK1 - cyclin B1 - MAD1 peptide complex have been deposited to the EM Data Resource with accession code EMD-100282 and the PDB with accession code 34ZF.

## Methods

### Cyclin B1 purification for phage display and fluorescence polarization

For phage display and fluorescence polarization experiments (FP), cyclin B1 and CDK2 were expressed in *E. coli* BL21 ArcticExpress DE3 cells (Agilent) as 6xHis-tagged proteins from pET28a-based vectors and purified using the Ni-NTA purification system (Thermo Fisher Scientific). The BL21 cultures were grown at 37°C to OD600 0.4, then grown in an 11°C shaker for 1h, followed by addition of 1 mM IPTG and overnight growth at 11°C. For cyclin B1, the cell pellets were lysed in a buffer containing 20 mM Tris-HCl pH 8.0, 800 mM NaCl, 10% glycerol, 10 mM imidazole, 0.5 mM TCEP-HCl, 1% Triton X100 supplemented with 1 mg/mL lysozyme and 1x cOmplete^TM^ protein inhibitors cocktail (Merck). CDK2 cell pellets were lysed in 50 mM Tris-HCl pH 7.4, 500 mM NaCl, 10% glycerol, 10 mM imidazole, 0.5 mM TCEP-HCl, 0.5% Triton X-100, 1 mg/mL lysozyme and 1x cOmplete^TM^ protein inhibitors cocktail (Merck) buffer. The His-tagged proteins were purified using Ni-NTA agarose and eluted with a buffer containing 200 mM imidazole. The eluates were then diluted 5x with 20 mM Tris-HCl pH 7.4, 300 mM NaCl, 0.1% Triton, 1 mM DTT buffer, concentrated using Amicon Ultra 4 10K centrifugal filters, divided into small aliquots and frozen in liquid nitrogen. For phage display, free cyclin B1 protein was used, while for FP experiments cyclin B1 and CDK2 were mixed in equimolar ratio.

### Phage display using the HD2 ProP-PD library

Phage selections using the HD2 ProP-PD library (∼950,000 overlapping 16-mer peptides from the intrinsically disordered regions of the human proteome) were carried out as described previously ^23^. Selections were performed in three sets of three replicates (two sets with Cyclin B1–Cdk2 and one with Cyclin B1 alone) over four consecutive rounds. In brief, target proteins (15 µg) or glutathione transferase (GST, control protein; 10 µg) were immobilized in 96-well MaxiSorp plates (Nunc) in a total volume of 100 µL diluted in phosphate-buffered saline (PBS) and the plates incubated overnight at 4°C. On day 1 of the selection, wells were blocked with 0.5% bovine serum albumin (BSA) in PBS (200 µL/well) and the plates incubated for 1 h at 4°C. The naïve phage library (10 µL/well, 10^11^ phages) was precipitated by adding one fourth of 20% PEG-8000 and 400 mM NaCl and incubation on ice for 15 min. The library was resuspended in 0.5% BSA in PBS and 0.05% Tween-20 (100 µL/well). The control plates with GST were washed four times with PBS supplemented with 0.05% Tween-20 and the naïve phage library was added to each coated well (100 µL). Plates were incubated for 1 h at 4°C. After that, plates with the immobilized target proteins were washed four times with PBS supplemented with 0.05% Tween-20 and the unbound phages were transferred from the GST plates to the respective wells of the protein plate. After 2 h incubation at 4°C, unbound phages from the protein plate were removed by washing five times with PBS supplemented with 0.05% Tween-20. Bound phages were eluted by adding log phase Omnimax bacteria and incubating for 30 min at 37°C and 200 rpm (100 µL/well). Next, M13K07 helper phages (10 µL/well, 10^9^ phages) were added to each well and the plates were incubated again for 45 min at 37°C at 200 rpm. The bacteria were subsequently amplified in 2YT medium supplemented with 30 µg/mL kanamycin, 100 µg/mL carbenicillin and 0.3 mM IPTG by incubation overnight at 37°C and 200 rpm. Proteins and GST were immobilized for the following selection round as described above. The steps on day 2-4 were the same as on day 1, with the exception that phages from the previous selection day were used as in-phages. For this, bacteria amplified overnight were spun down for 15 min at 1,700 xg and 4°C. The supernatant containing the phages was pH- neutralized by the addition of 10x PBS and subsequently heat-inactivated by incubation at 60°C for 10 min. Phages harvested in this fashion were then added to the respective well of the GST control plate (100 µL). Phage pools were stored at 4°C and additionally used in phage pool ELISA (enzyme-linked immunosorbent assay) and next generation sequencing (NGS).

### Phage pool ELISA

To assess enrichment of the phage pools for binders to the proteins, phage pool ELISA was performed as previously described ^23^. For this, proteins (7.5 µg) and GST (5 µg) were immobilized on a 384-well MaxiSorp plate (Nunc) (total of 50 µL) and the plates were incubated overnight at 4°C. Plates were blocked for 1 h with 100 µL 0.5% BSA in PBS for 1 h at 4°C. Phage pools from selection day 1-4 were added to both target protein and GST wells and the plates incubated for 1.5 h at 4°C. After incubation, plates were washed four times with PBS supplemented with 0.05% Tween-20 (100 µL/well) and bound phages were detected by the addition of anti-M13 mouse HRP-conjugated antibody (50 µL/well, 1:5000 in 0.5% BSA in PBS supplemented with 0.05% Tween-20, Sino Biological Inc). Plates were incubated for 1 h and then washed four times with PBS supplemented with 0.05% Tween-20 and once with PBS (100 µL/well). After that, 40 µL of TMB substrate (Seracare KPL) were added to each well, which was converted by the HRP if there was a bound antibody. The reaction was stopped by the addition of 0.6 M H_2_SO_4_ (40 µL/well) and the signal was read out on an iD5 (Molecular Devices) by measuring the absorption at 450 nm. Pools were considered enriched if the signal of the protein was at least two-fold higher in comparison to control wells.

### Next generation sequencing and analysis pipeline

The peptide coding regions of the enriched phage pool were barcoded and PCR amplified using a dual barcoding strategy ^23^. For this purpose, 5 µL of each phage pool were used as template in a PCR reaction of 22 cycles using Phusion High-Fidelity PCR Master Mix (Thermo Fisher). PCR reactions (25 µL) were normalized by purification with 25 µL Mag-bind Total Pure NGS (Omega Bio-tek). The amplicons were pooled and further purified by gel purification in a 1% agarose gel supplemented with GelRed (1:10000) using the QIAquick Gel extraction Kit Qiagen). DNA content was quantified with a Quant-iT PicoGreen dsDNA Assay Kit (Molecular probes by Life technologies) and the sample was sent for NGS to the NGS-NGI SciLifeLab facility. The samples were analyzed by Illumina MiSeq v3 run, 1 × 150 bp read setup, 20% PhiX and returned an average of 33,000 reads per set of barcodes (on average 18 million reads per NGS run).

### Fluorescence polarization

N-terminally FITC-Ahx labeled PKN2-RxL peptide (SAVRKLDFSD) and MAD1 helix peptide (GSLQMQYQQSMQLEERA) were ordered from GenScript with >95% purity. Fluorescence polarization was measured in a buffer containing 25 mM Hepes-KOH pH 7.4, 150 mM NaCl, 5 mM DTT and 0.05% CHAPS. The assays were carried out in black, low volume, non-binding, 384-well plates (4514, Corning). After mixing, the reactions were incubated at 37°C in the dark for 30 minutes before measuring fluorescence polarization using the PHERAstar FSX microplate reader (BMG Labtech). The fluorescence polarization experiments were performed in at least two replicates. The data was analyzed and plotted using Prism (GraphPad).

### Cyclin B1 purification for Cryo-EM

CDK1 and cyclin B1 were expressed in High5 (*Trichoplusia ni*) cells using bacmids generated from pLIB-GST-3C-CDK1 (#177011, Addgene), pLIB-HIS-TEV-scCAK1 (#177014, Addgene), and pLIB-His6-TEV-cyclin B1 (#177012, Addgene) plasmids deposited by A. Musacchio ^41^. Cyclin B1 was purified by nickel affinity chromatography followed by cleavage by tobacco etch virus (TEV) protease and size exclusion chromatography; phosphorylated CDK1 was purified by GST-affinity chromatography, cleavage by PreScission protease, and size exclusion chromatography. The cyclin B1-CDK1 complex was formed by incubation of the separately-purified proteins on ice followed by size exclusion chromatography ^42^.

### Cryo-EM Sample preparation

Cryo-EM samples were prepared on UltrAuFoil R 1.2/1.3 300 mesh holey gold grids (Quantifoil Micro Tools). Samples were prepared by mixing the Cyclin B1-CDK1 complex (5x dilution from 3.3 mg/mL stock) with 200 mM of MAD1 peptide (20x dilution from 10 mg/mL stock) in cryo-EM buffer (20 mM HEPES-NaOH pH 7.5, 250 mM NaCl). 4 mL of each complex were applied to plasma-cleaned grids (Tergeo EM plasma cleaner, PIE Scientific), which were blotted for 1.5-2.0 s using a Vitrobot Mark IV (Thermo Fisher Scientific) operated at 5°C and 100%[Nd1] humidity, and then plunged-frozen into liquid ethane cooled by liquid nitrogen. After vitrification, the grids were clipped into AutoGrid cartridges (Thermo Fisher Scientific) to use with autoloader systems.

### Data acquisition

Grids were loaded into a Titan Krios G3i cryo-transmission electron microscope (cryo-TEM) (Thermo Fisher Scientific) operated at 300kV acceleration voltage and equipped with a K3 direct electron detector (Gatan) and a BioQuantum energy filter (Gatan). Data were collected as dose-fractionated movies (70 frames) in TIFF format at 165,000x nominal magnification using 2x hardware binning (counted super-resolution bin 2), resulting in a pixel size of 0.51 Å/pixel, with a total exposure of 70 electrons/Å^2^, and a defocus range of -0.6 to -1.8 µm. Grid squares were selected manually and holes for data acquisition were automatically detected and selected using appropriate relative ice thickness filters in EPU Software (Thermo Fisher Scientific) for Cyclin B1-CDK1 complexes. In total, 14,745 movies were collected from one grid.

### Cryo-EM image processing

Cryo-EM data were pre-processed in CryoSPARC live and CryoSPARC v4.4.1 ^43^, and then transferred to RELION 5.0.0 ^44^ for further processing, 3D reconstruction, and final refinement. All refinements were performed using BLUSH regularization implemented in RELION 5.0.0 ^44^.

In CryoSPARC live, raw movies were motion corrected with 2x binning. The contrast transfer function (CTF) of each motion-corrected micrograph was fitted in CryoSPARC live. Total beam-induced specimen motion, in-frame motion, relative ice-thickness, and the quality of CTF fitting were inspected to remove poor-quality micrographs, which led to 7,873 micrographs being accepted for Cyclin B1-CDK1-MAD1. Particles were picked using blob picker (circular/elliptical blob, 70-90 Å diameter) and then extracted using a box size of 160x160 pixels. On-the-fly 2D classification was used to assess data quality.

To maximize the selection of high-quality particles, two particle selection strategies were employed in CryoSPARC v4.4.1 ^43^ after completion of on-the-fly processing. All blob-picked particles from CryoSPARC live session were re-extracted using a box size of 80x80 pixels and re-classified into 100 2D classes using a batch size of 300 particles, which resulted in 611,761 particles retained (particle set (i) identified in Supplementary Figure S1). Particles from good 2D classes from the CryoSPARC live session were re-extracted using a box size of 80x80 pixels and re-classified in 100 2D classes using a batch size of 300 particles which resulted in 450,150 retained particles retained (particle set (ii) from Supplementary Figure S1). Particles were also extracted using template picking using 2D averages from CryoSPARC live. Template-based picked particles were classified into 100 2D classes using a batch size of 300 particles which resulted in 595,004 retained particles from good 2D classes retained (particle set (iii) identified in Supplementary Figure S1). All particle subsets (particles (i)-(iii)) were combined and duplicates were removed, resulting in 955,005 retained particles. These particles were then classified into 50 2D classes using a batch size of 200, resulting in 801,099 retained particles, which were then converted to *.star files for use in RELION 5.0.0 using scripts contained in the PYEM package ^45^.

Motion-corrected micrographs from CryoSPARC live were imported to RELION 5.0.0 and selected particles were re-extracted using a box size of 160x160 pixels using the coordinate information from the imported particles from CryoSPARC v4.4.1. Extracted particles were subjected to masked 3D auto-refinement using a cryo-EM structure of CDK2-cyclin A as the initial reference ^46^. Particles were then classified into four 3D classes by 3D classification without alignment (regularisation parameter τ = 20). A total of 146,618 particles were selected from the best 3D class and refined to 3.4 Å resolution. To further improve the map quality, selected particles were subjected to CTF refinement (beam tilt and trefoil). CTF-refined particles were then re-refined and classified into three 3D classes by 3D classification without alignment (regularization parameter τ = 20). Finally, 57,945 particles were selected from the best 3D class and were refined and post-processed yielding a 3.3 Å reconstruction of the Cyclin B1-CDK1-MAD1 complex, according to Fourier shell correlation (FSC) at the 0.143 cut-off (Supplementary Figure S2A).

### Cryo-EM model building and refinement

To build the Cyclin B1-CDK1-MAD1 structure, an Alphafold-predicted model ^30^ was fitted into the post-processed cryo-EM map (3.3 Å resolution) using UCSF Chimera ^47^. The fitted model was transferred to Coot ^48^ for model building and refined by real-space refinement in PHENIX ^49^. The refined model was validated using MolProbity ^50^ implemented in PHENIX. Refinement statistics are provided in Supplementary Table S1. The 3D Fourier shell correlation (FSC) of the reconstruction was computed using the remote 3DFSC Processing server ^51^.

### Cryo-EM visualization of molecular models and creation of figures

Molecular models and cryo-EM maps were visualized in UCSF Chimera ^47^, UCSF Chimera X ^52^ and PyMOL (Schrödinger, Inc.) for analysis, interpretation and figure preparation.

### Extraction of hydrophobic patch binding peptides from Predictome models

We retrieved the PDB files of all human cyclin B1 and B2 complexes with non-zero SPOC scores from Predictome ^32^, obtaining 127 and 111 complexes of cyclin B1 and B2, respectively, with each interactor pair represented by three PDB models. We extracted the peptide sequence that interacts with the cyclin hydrophobic patch, defined by the peptide residues being less than 4 Å from positions 200-212 in cyclin B1 and 168-180 in cyclin B2. The peptides were elongated into 16mers, resulting in 327 unique peptides.

### Cloning and yeast strains for SIMBA

The yeast strains used in this study are in the common laboratory strain background W303 and are described in Supplementary Table S4. Standard methods were used for growth and genetic engineering of yeast. Yeast were grown at 30°C in yeast extract/peptone medium with 2% glucose (YPD) or in synthetic complete medium (SC) lacking uracil (or uracil and histidine) supplemented with 2% glucose or raffinose. The human cyclin reading frames were cloned between GST and yeast CLN2 in a *P_GAL1_-GST-CLN2* expression cassette, pPP4343 or pPP3572 ^17,24^. As described previously ^17^, the intrinsically disordered N- and/or C-termini of the human cyclins were truncated to remove localization and degradation signals, and a triple alanine mutation was introduced in the conserved CDK-binding interface of each cyclin to prevent the human cyclins from interacting with yeast Cdk1. These cyclin expression constructs were harbored on replicating (CEN/ARS) plasmids or were integrated at the HIS3 locus in yeast (Supplementary Table S4). The cyclin docking peptides, flanked by GSGG flexible linker sequences, were inserted into a Ste20^Ste5PM^chimera and expressed in yeast from a low copy number plasmid with URA3 selection. Oligonucleotide pools encoding docking peptides were ordered from GenScript and were amplified by PCR (12 cycles) using primers that anneal to the common flanking linkers and include MluI and SphI restriction sites. For deep mutational scanning of MAD1, template oligonucleotides were designed such that single codons were randomized (NNN) to generate all 64 nucleotide variants, and then these were amplified by PCR as above. The PCR products were digested with MluI and SphI, treated with calf intestinal phosphatase, and then ligated into the MluI/SphI-digested Ste20^Ste5PM^ expression vector (pPP4745). The ligation products were transformed into *E. coli* (XL-10 Gold Ultracompetent Cells; Agilent/Agilent Technologies) and plated on LB+Amp plates.

Two libraries were prepared and screened independently. Library 1 included 304 MAD1 mutational scanning variants, 327 candidate B1/B2-binding peptides from the Predictome database, 8 MAD1 orthologs from non-human vertebrates, and 51 RxL peptides (44 wild-type plus 7 single-position variants). Library 2 included 43 peptides matching the loose MAD1-like consensus sequence, 4 MAD1 variants (2 non-binders, 2 optimized), and 43 wild-type RxL peptides. Both libraries contained negative controls (known non-binding peptides) and positive controls (Cln2-binding LP peptides and STOP codon peptides). All peptides were encoded by three synonymous oligonucleotides with the exception of the MAD1 mutational scanning variants, which were generated by codon randomization (NNN) and hence were represented by one to six oligonucleotides each.

### SIMBA signaling assays

SIMBA is a yeast-based assay of domain-peptide binding ^24^, in which bait domains are fused to the yeast cyclin Cln2 and peptides are inserted in a chimeric signaling protein Ste20^Ste5PM^; then, binding of the bait domain to the peptide allows Cln2-Cdk1 mediated phosphorylation of Ste20^Ste5PM^, which inhibits signaling in the yeast mating pathway. For low-throughput assays of cyclin binding to the Ste20^Ste5PM^chimera, mating pathway signaling was measured by a *P_FUS1_-lacZ* transcriptional reporter construct driving β-galactosidase expression, as described previously ^17,24^. For this, yeast cultures were grown in SC medium containing raffinose and, at OD600 0.7, the exogenous cyclin expression was induced by adding galactose (2% final). After 90 min, mating signaling was activated by adding α-factor to 50 nM and the cultures were grown for an additional 45 min. Then, the yeast cells were pelleted and resuspended in 60 mM disodium phosphate, 40 mM monosodium phosphate, 10 mM KCl and 1mM magnesium sulfate buffer. The cells were permeabilized by addition of chloroform (to 10%) and sodium dodecyl sulfate (to 0.008%), and then ortho-nitrophenyl-β-galactoside was added (to 0.9 mg/mL) as a spectrophotometric substrate for β-galactosidase activity. The reactions were incubated at 30°C. β-galactosidase activity was estimated by measuring formation of ortho-nitrophenol in the reaction supernatant at OD420.

### SIMBA competitive growth experiments

For high-throughput assays of cyclin-peptide binding, pooled Ste20^Ste5PM^ expression plasmid libraries were transformed into yeast strains harboring integrated expression cassettes for different HsCyclin-Cln2 fusions, as described previously ^17^. The transformation mixture was plated on SC medium plates lacking uracil and incubated at 30°C for 3 d. Diluted aliquots from the transformation were used for colony counting to ensure that the number of transformants exceeded the number of nucleotide sequences in the library by at least 10-fold. The transformed cells were collected from the plates, diluted into 50 mL SC-URA containing raffinose and grown at 30°C for 4–8 h. Then, the 50-mL cultures were diluted to OD660 0.05 and grown overnight. At OD660 0.8–1, expression of the bait domain-Cln2 fusion was induced by adding 2% galactose. After 75 min, an aliquot (∼3 × 10^8^ cells) was collected from each culture via centrifugation and flash frozen in liquid nitrogen. At the same time, yeast mating pheromone (α-factor) at 500 nM final concentration was added to the culture to begin the competitive growth. The cultures were grown at 30°C and diluted every 12 h to keep the OD660 below 1. Aliquots (20 mL; ∼3 × 10^8^ cells) were harvested at 0, 20 and 32 h after α-factor addition (t0, t20, t32). Harvested cells were washed with water, transferred to 1.5-mL tubes and centrifuged again. The resulting cell pellets were flash frozen in liquid nitrogen and stored at -80°C.

### Preparation of SIMBA samples for deep sequencing

As described previously ^17,24^, DNA was purified from yeast cells using a ZR Plasmid Miniprep Kit (Zymo Research, cat. no. D4015). Frozen cell pellets were thawed and suspended in 200 μL solution P1 before they were lysed using Zymolyase (0.2 units/μL; Zymo Research, cat. no. E1005) for 1.5 h at 37°C. Plasmid DNA samples (4 μL) were subjected to PCR (17 cycles, 50 μL total volume) with primers that include standard P5 and P7 sequences for binding to Illumina flow cells during NGS. The forward primer included a P5 sequence followed by an Illumina sequencing primer binding site, a 6-nucleotide bar code and a plasmid-annealing sequence upstream of the peptide-encoding insert; the reverse primer included a P7 sequence followed by a 6-nucleotide i7-index sequence, an i7 sequencing primer binding site and a plasmid-annealing sequence downstream of the peptide-encoding insert. Aliquots (5 μL) of the PCR products were run in 1.2% agarose gels to confirm the presence of the desired product, and the remainders were purified using Zymo-Spin I Columns (Zymo Research, cat. no. C1003-250) and eluted in 10 mM Tris-HCl, pH 8.0. The concentration of the eluted products was measured using a NanoDrop ND-1000 (Thermo Fisher Scientific), and a mixture was prepared containing equal amounts of each DNA product. The PCR amplicon libraries were sequenced by Novogene USA (Sacramento, CA).

### SIMBA data analysis

2FAST2Q ^53^ was used to obtain the counts of different peptide-encoding oligonucleotide sequences from the deep sequencing data. To calculate enrichment scores, the following data processing steps were performed: (1) oligonucleotides with <20 reads at t0 were dropped from further analysis. (2) The frequency of each oligonucleotide within a sample at t0, t20 and t32 was calculated. (3) The log2 fold change of t20 and t32 frequencies compared to the t0 frequency was calculated, and then divided by the timepoint (in hours), resulting in a log2 slope value. (4) The log2 slope values were averaged for t20 and t32. (5) For each bait experiment, the average log2 slope values were z-normalized using the mean and standard deviation of a set of negative controls. (6) For each peptide, the median z-score in the unfused Cln2 strain was subtracted from the median z-score in each bait domain Cln2 fusion strain to obtain the final SIMBA scores. For most peptides, these median values were derived from six measurements (three synonymous oligos in two replicate experiments). For the mutant MAD1 variant peptides in DMS experiments, which were generated by random codon mutagenesis (NNN) that creates different numbers of codons for each amino acid, the median values were derived from two to twelve measurements of each mutant variant (one to six synonymous oligos in two replicate experiments) plus 80 measurements of wild-type MAD1 (40 synonymous oligos in two replicate experiments). To create logos of sequence preferences derived from DMS data, SIMBA scores were transformed into a difference matrix ^54^. First, the score for each amino acid variant was normalized to the lowest and highest scores of all variants. Then, all normalized values at a given residue position were adjusted by subtracting the average normalized value at that same position. Logos were plotted from the adjusted matrix using the Python package ‘Logomaker’ ^55^.

### Cell culture

HEK293T and RPE1-hTERT-FRT cells were cultured in high glucose DMEM (41966029, Gibco) supplemented with 110 mg/L pyruvate, 3.7 g/L sodium bicarbonate, 10% fetal bovine serum (A5209402, Gibco), 100 U/mL penicillin, and 0.1 mg/mL streptomycin (15140122, Gibco) in a 37°C incubator with 5% CO_2_. All cell lines were tested negative for mycoplasma by qPCR at the Cancer Research UK Cambridge Institute Research Instrumentation and Cell Services core facility (RRID:SCR_028062). Doxycycline hyclate (D9891-1G, Sigma-Aldrich) was dissolved at 1 mg/mL in DMSO. Palbociclib (S1116, Selleck) was dissolved at 10 mM in DMSO and nocodazole (13857, Cayman Chemical) at 2 mg/mL DMSO.

### Generating CDK reporter expression cell lines

CDK activity reporters were based on the intrinsically disordered C terminus of PKMYT1, where the native RxL motif was replaced with different cyclin docking motifs ^17^. The CDK reporters were fused to a 3x FLAG tag, EGFP, and a nuclear localization signal, and expressed from a doxycycline-inducible lentiviral vector (pCW57.1, Addgene #41393). The virus was produced as follows: 7 x 10^5^ HEK293T/17 cells were seeded to 6-well plate, and after 24 hours were transfected using lipofectamine 3000 (Thermo Fisher Scientific) with psPax2 (Addgene #12260) and pMD2.G (Addgene #12259) as packaging plasmids. Six hours after the transfection, the medium was replaced with 3 ml fresh DMEM. The medium containing the virus was collected 2 days after transfection and was used to transduce RPE1-FRT cells. 24 hours before transduction, 2 x 10^5^ RPE1-hTERT-FRT cells were seeded to a 6-well plate. For transduction, the RPE1 cell media was replaced with the lentiviral supernatant and supplemented with 8 µg/mL polybrene, followed by culturing the cells for 2 days. Then, the transduced RPE1 cells were collected by trypsinizing and seeded to a 6 cm dish in DMEM supplemented with 1 µg/mL puromycin. Prior to seeding for experiments, the transduced RPE1 cells were split at least 3 times and cultured in the presence of puromycin to allow selection of the transduced cells.

### Co-immunoprecipitation

For CDK reporter co-immunoprecipitation experiments, the RPE1-based cell lines were seeded to 15 cm dishes at 40% confluency, 1 µg/mL doxycycline was added to the medium to induce the reporter expression and the cells were collected and frozen at 80% confluency 24 hours after induction. The cells were lysed in a buffer containing 10 mM Tris-HCl pH 7.4, 150 mM NaCl, 0.1% NP-40, 5 mM EDTA, 1 mM DTT and cOmpleteTM protein inhibitors cocktail (Merck). The total protein concentration in the lysate was measured using Pierce™ Bradford Plus Protein Assay Reagent (Thermo Fisher Scientific). The lysates were diluted to 2 mg/mL total protein. For co-immunoprecipitation, 500 µg of cell lysate was mixed with 10 µL ChromoTek GFP-Trap® Magnetic Agarose beads (Proteintech) that had been equilibrated with the lysis buffer. The mixture was incubated on a rotator at 4°C for 1 hour. Then, the GFP-trap magnetic beads were collected using a magnet, the lysate was removed and the beads were washed three times with 500 µL of lysis buffer. The immunoprecipitated proteins were eluted using Laemmli SDS-PAGE sample buffer and were subjected to SDS-PAGE and immunoblotting.

### Immunoblotting

To analyze the phosphorylation of the CDK reporters in the cell cycle, the synchronized RPE1 cells expressing the reporter-GFP fusion proteins were lysed in a buffer containing 10 mM Tris-HCl pH 7.4, 150 mM NaCl, 1% Triton X100, phosSTOP phosphatase inhibitor cocktail (Roche) and cOmplete EDTA-free protease inhibitor cocktail (Merck). The lysate was cleared by centrifugation and the total protein concentration was measured using Pierce™ Bradford Plus Protein Assay Reagent (Thermo Fisher Scientific). For immunoblotting, 20 µg of cell lysate was resolved using Phos-tag SDS-PAGE with gels containing 8% acrylamide, 25 µM Phos-tag (Alpha Laboratories) and 50 µM MnCl_2_ as previously described ^56^. Following electrophoresis, the gels were washed twice with 10 mM EDTA. The proteins were transferred to the nitrocellulose membrane using iBlot 3 Broad Range preprogrammed method (Thermo Fisher Scientific). After transfer, the membrane was blocked using 5% fat-free milk solution in Tris-buffered saline with Tween-20 (TBS-T), followed by overnight incubation at 4°C with the primary antibody solutions. The following primary antibodies were used: sc-271682 (Santa Cruz Biotechnologies) at 1:500 for cyclin A, sc-245 (Santa Cruz Biotechnologies) at 1:500 for cyclin B1, 21644-1-AP (Proteintech) at 1:1000 for cyclin B2, XP® Rabbit mAb #5174 (D16H11) (Cell Signaling Technologies) at 1:1000 for GAPDH, D6W5B Rabbit mAb (Cell Signaling Technologies) at 1:1000 for FLAG tag. Following primary antibody incubation, the membrane was washed 5 times with TBS-T, incubated with secondary antibody solutions for 1 hour, and washed again 5 times with TBS-T. HRP-conjugated anti-mouse IgG (#7076, Cell Signaling Technologies) at 1:2000 and HRP-conjugated anti-rabbit IgG (#7074, Cell Signaling Technologies) at 1:10000 were used as secondary antibodies. The antibodies were detected using SuperSignal™ West Pico PLUS Pico Chemiluminescent substrate (Thermo Fisher Scientific).

### RPE1 cell cycle synchronization

For cell cycle synchronization, 2.5 x 10^5^ RPE1-FRT cells containing the CDK reporter expression cassettes were seeded to 10 cm dishes and grown for 32 hours. Then, the medium was replaced with DMEM containing 150 nM palbociclib to arrest cells in G1 phase and 1 µg/mL doxycycline to induce the expression of CDK reporters. The cultures were incubated at 37°C for 24 hours, after which the palbociclib-containing medium was removed, the cells were washed 3 times with 10 mL DMEM and the cells were grown in DMEM supplemented with 1 µg/mL doxycycline at 37°C. At 0, 4, 6, 10 and 14 hours, the cells were collected by trypsinization, washed with Tris buffered saline and flash frozen for immunoblotting. For propidium iodide staining, the cells were fixed with 70% ethanol and stored at -20°C.

### Propidium iodide staining and flow cytometry

The ethanol-fixed RPE1 cells were washed twice with PBS and resuspended in 300 µl solution containing 5 µg/mL propidium iodide (421301, Biolegend) and 25 µg/mL RNase A in PBS. The mixture was incubated at 37°C in the dark for 30 minutes and analyzed by flow cytometry using BD LSR Fortessa. Data from at least 10 000 cells was collected and analyzed using FlowJo (BD Biosciences).

## Supplementary Figures

**Supplementary Figure S1:**
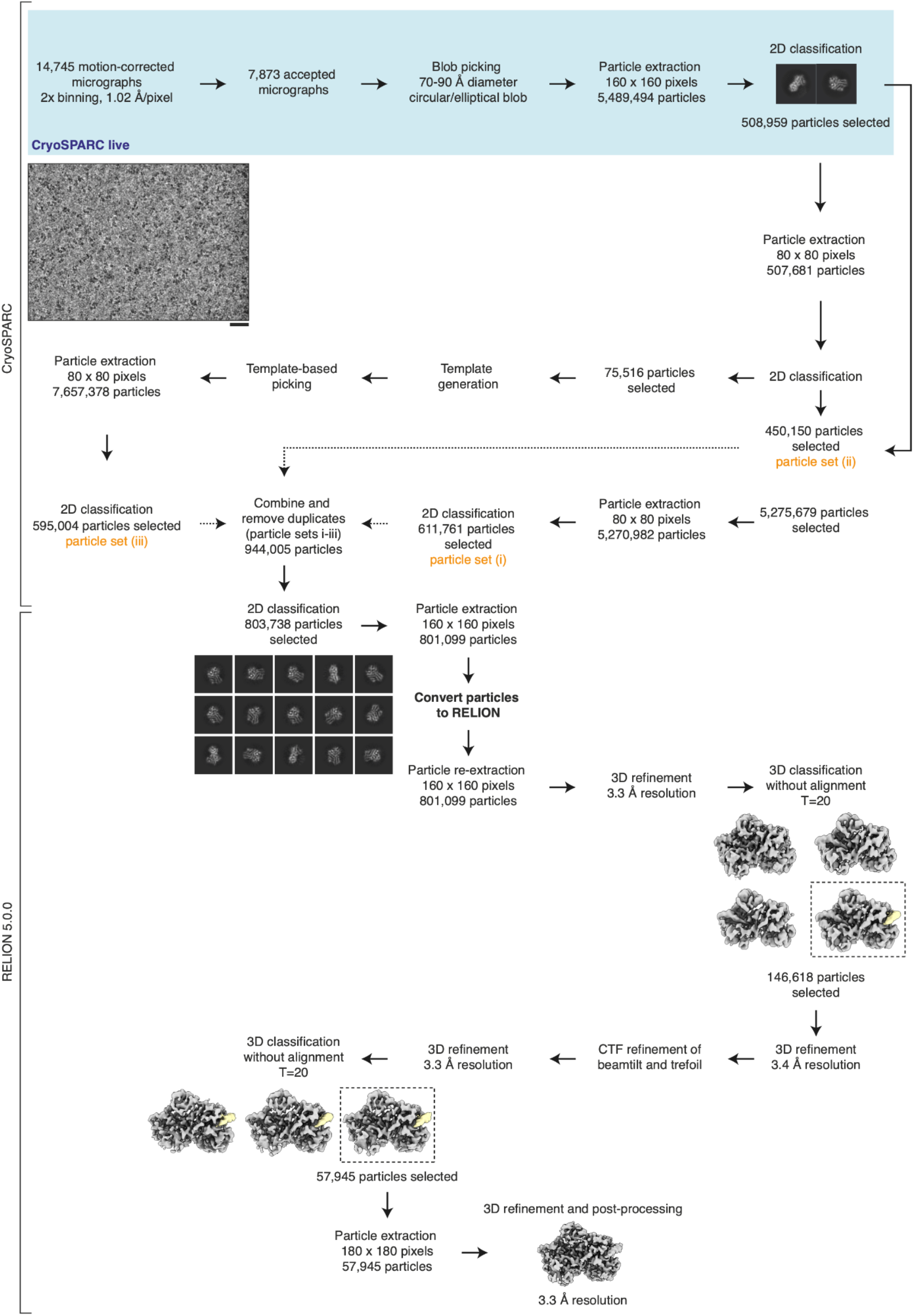
Cryo-EM data processing for the Cyclin B1-CDK1-MAD1 reconstruction. Cryo-EM processing workflow used to obtain the reconstruction of the Cyclin B1-CDK1-MAD1 complex. A representative micrograph is shown (scale bar: 25 nm). The micrograph was low-pass filtered to 8 Å. 3D refined maps and maps from 3D classification are shown as surface views with the MAD1 helix highlighted in pale yellow. Classes which are taken forward for further processing are indicated with dashed-line boxes.

**Supplementary Figure S2:**
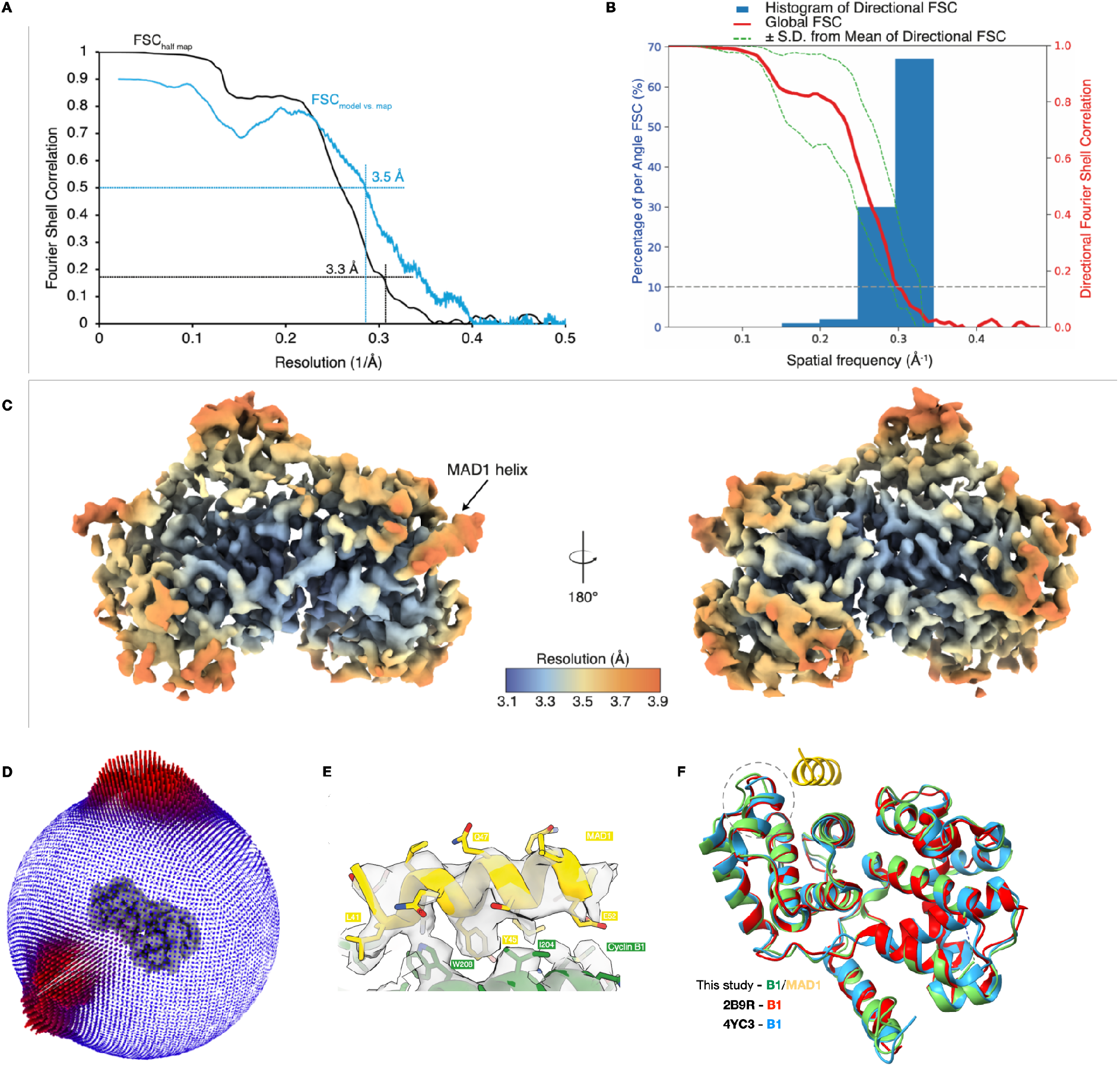
Overall and local resolutions of the Cyclin B1-CDK1-MAD1 reconstruction. **(A)** Gold standard (black) and model vs. map (blue) Fourier shell correlation (FSC) plot evaluated at the appropriate thresholds. **(B)** Histogram of directional FSCs plots produced by the 3D FSC validation server ^51^. The sphericity of 3D FSC of the reconstruction is 0.81. **(C)** Results from local resolution estimation computed in RELION. **(D)** Orientation distribution plot from RELION 3D auto-refinement. **(E)** Cryo-EM density of the MAD1 peptide. **(F)** Structure overlay of the MAD1 peptide (yellow) bound structure of Cyclin B1 (green) with two apo-structures (PDB: 2B9R (red) and PDB: 4YC3 (blue)). Conformational shift of residues 271-273 at the C-terminus of the α4 helix highlighted by a dashed circle.

**Supplementary Figure S3:**
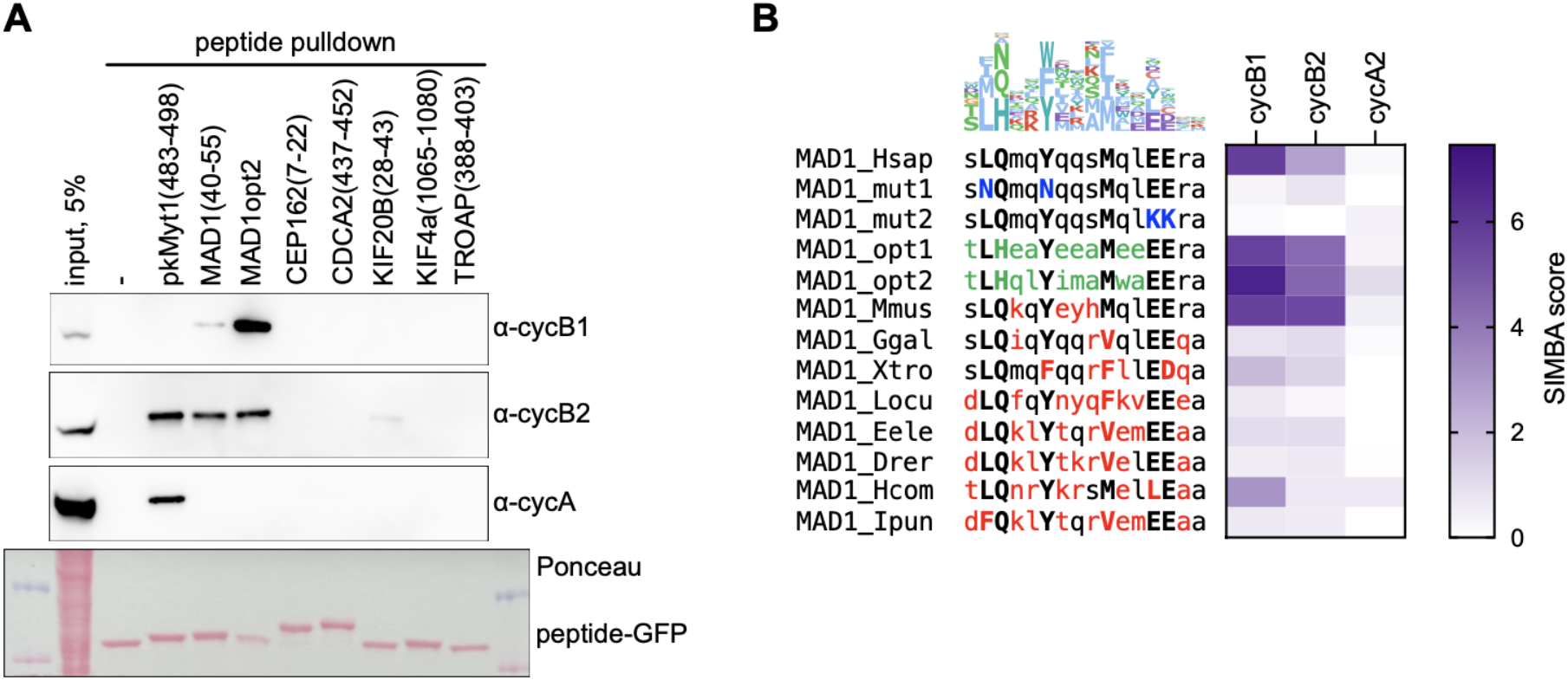
Analysis of additional candidate MAD1-like peptides. (**A**) Peptide pulldown assay testing five Predictome hits for the ability to bind B- or A-type cyclins in HCT116 cell extracts. Peptides were expressed as fusions to GFP. Controls include no peptide (-), an RxL peptide from PKMYT1, plus wild-type and opt2 versions of the MAD1 peptide. A representative example of two independent experiments is shown. (**B**) SIMBA scores for binding of cyclins B1, B2, and A2 to MAD1 orthologs from eight non-human vertebrates. Control peptides include wild-type human MAD1 (MAD1_Hsap) and its non-binding and optimized variants defined in Figure 3D, with mutations denoted by blue or green font. Red font denotes residues in orthologs that differ from human MAD1. Uppercase, cyclin B1 contacting positions; lowercase, non-contacting positions. The logo at top shows residues preferred by cyclin B1 determined from DMS data.

**Supplementary Figure S4:**
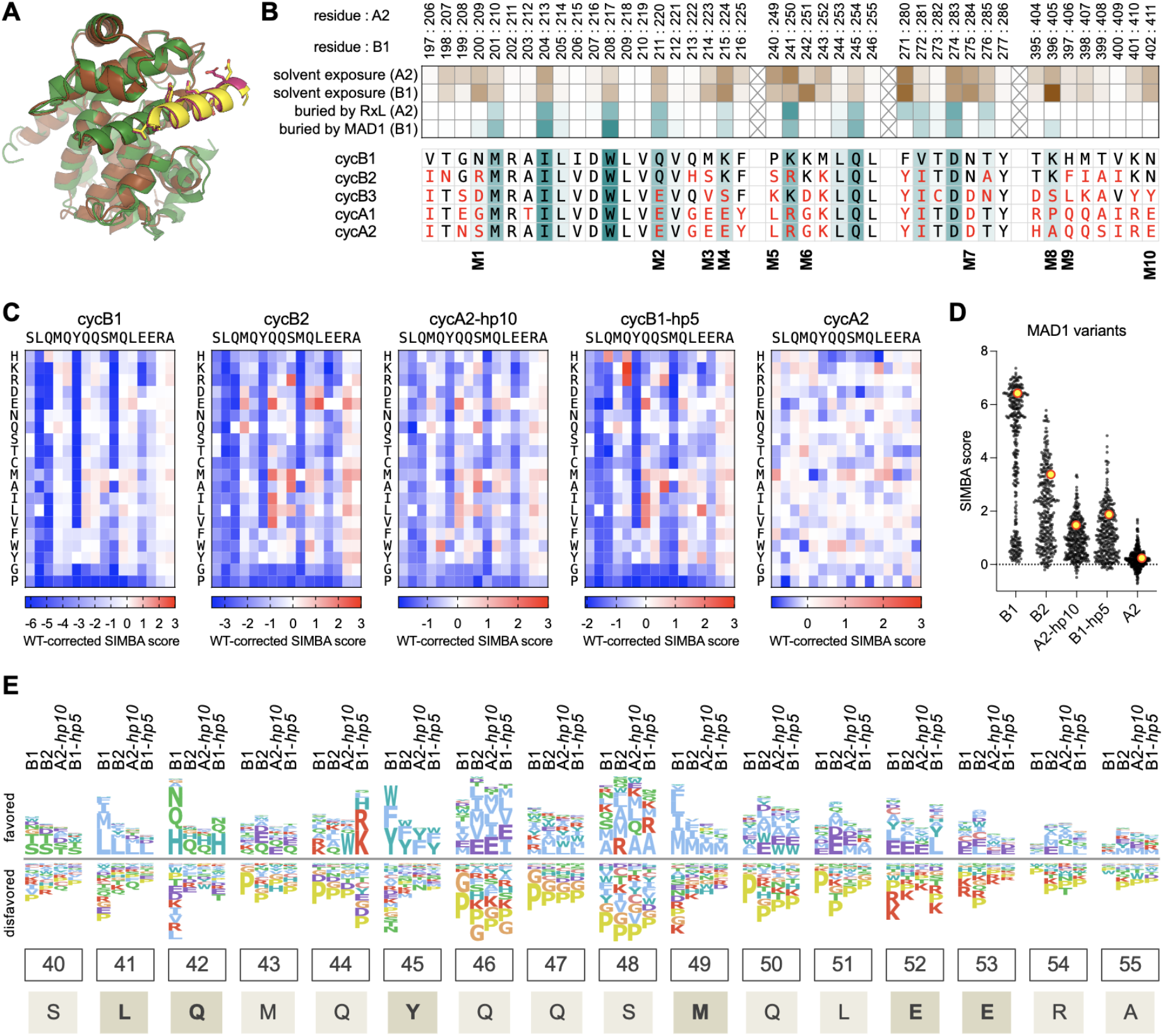
Comparison of Cyclins B1, A2, and their pocket-swapped derivatives. **(A)** AlphaFold3 models of MAD1 (yellow) bound to Cyclin B1 (green; ipTM = 0.95, pTM = 0.94), or MAD1 (pink) bound to Cyclin A2 (brown; ipTM = 0.96, pTM = 0.96). **(B)** Cyclin residues in peptide-binding pockets. Top, for residue in cyclins A2 and B1, heatmaps show the relative levels (in Å²) of solvent exposure (in the unbound state) and buried surface area (in the bound state), as calculated from the A2-E2F1 complex (PDB ID: 1H24) or the B1-MAD1 complex (this study). Bottom, comparison of residues in five Cyclins (B1, B2, B3, A1, A2), shaded according to B1-MAD1 burial, with differences from B1 marked in red. Positions mutated in the pocket swap analyses of Figure 4 are indicated below (M1 – M10). **(C)** Heatmaps showing MAD1 DMS results for the indicated cyclins and pocket-swap derivatives. **(D)** SIMBA score distribution of all MAD1 DMS variants for each cyclin. Large yellow circles, wild-type MAD1; small black circles, mutant variants. **(E)** Logo derived from MAD1 DMS data comparing residue preferences of four cyclins at each MAD1 peptide position. All cyclins share similar preferences at the core contact residues. At non-core positions, notable differences include: (i) at position 44, B1 and B1-*hp5* strongly prefer Arg/Lys, whereas B2 and A2-*hp10* do not and instead A2-*hp10* strongly prefers Trp; (ii) at position 46, all domains prefer aliphatics or Glu, but not Asp; (iii) at position 48, B2 and B1-*hp5* show a unique preference for Arg; and (iv) at positions 51 and 52, B1-*hp5* lacks the preference for acidic side chains shown by other domains.

## Supplementary Tables

**Table S1:** Cryo-EM data collection, refinement, and validation statistics.

| Dataset | Cyclin B1-CDK1- MAD1 |
|---|---|
| Microscope | Titan Krios G3i |
| Stage type | Autoloader |
| Acceleration voltage (kV) | 300 |
| Detector | Gatan K3 |
| Energy filter | Gatan Bio Quantum |
| Acquisition mode | 2x hardware binning |
| Pixel size (Å) | 0.51 |
| Defocus range (µm underfocus) | 0.6-1.8 |
| Electron exposure (e<sup>-</sup>/A<sup>2</sup>) | 70 |
| **Reconstruction** | EMD-100282 |
| Software | RELION 5.0.0 |
| Particles after 2D classification | 1,938,556 |
| Particles final | 57,945 |
| Extraction box size (pixels) | 160 x 160 x 160 |
| Final pixel size (Å) | 1.02 |
| Accuracy rotations (°) | 2.45 |
| Accuracy translations (Å) | 0.64 |
| Map resolution (Å) | 3.3 |
| Map resolution range (Å) | 3.1-4.1 |
| Sphericity | 0.812 |
| Map sharpening B-factor (Å<sup>2</sup>) | -50 |
| **Coordinate refinement** |  |
| Software and algorithm | PHENIX (real space refine) |
| Resolution cutoff (Å) | 3.3 |
| FSC<sub>model-vs-map</sub>=0.5 (Å) | 3.5 |
| **Model** | PDB-34ZF |
| Number of residues | 526 |
| Protein | 526 |
| B-factors overall | 78.06 |
| R.M.S. deviations |  |
| Bond lengths (Å) | 0.003 |
| Bond angles (°) | 0.589 |
| **Validation** |  |
| Molprobity score | 1.99 |
| Molprobity clashscore | 9.25 |
| Rotamer outliers (%) | 1.69 |
| C<sub>β</sub> deviations (%) | 0.0 |
| Ramachandran plot |  |
| Favored (%) | 96.27 |
| Allowed (%) | 3.73 |
| Outliers (%) | 0.0 |
| Rama-Z scores |  |
| Whole | 1.50 |
| Helix | 1.94 |
| Sheet | -1.25 |
| Loop | 0.04 |

**Supplementary Table S2.** High-throughput SIMBA scores of peptides matching the MAD1 consensus sequence.

**Supplementary Table S3.** High-throughput SIMBA scores of Predictome peptides and MAD1 DMS variants.

**Supplementary Table S4.** Yeast strains, plasmids, oligos for SIMBA experiments.

